# A Novel Performance Metric for Multiclass Subject Invariant Brain Computer Interfaces with Imbalanced Classes

**DOI:** 10.1101/2020.02.07.938548

**Authors:** Jesse Sherwood, Jesse Lowe, Reza Derakhshani

**Affiliations:** Computational Intelligence and Bio-Identification Technologies (CIBIT) Lab, University of Missouri – Kansas City, MO 64110, USA

**Keywords:** Brain Computer Interface, Machine Learning, Support Vector Machines, Human Movements, Multiclass Discriminants, Sequential Floating Forward Selection, Imbalanced Classes

## Abstract

[Finding suitable common feature sets for use in multiclass subject independent brain-computer interface (BCI) classifiers is problematic due to characteristically large inter-subject variation of electroencephalographic signatures. We propose a wrapper search method using a one versus the rest discrete output classifier. Obtaining and evaluating the quality of feature sets requires the development of appropriate classifier metrics. A one versus the rest classifier must be evaluated by a scalar performance metric that provides feedback for the feature search algorithm. However, the one versus the rest discrete classifier is prone to settling into degenerate states for difficult discrimination problems. The chance of occurrence of degeneracy increases with the number of classes, number of subjects and imbalance between the number of samples in the majority and minority classes. This paper proposes a scalar Quality (Q)-factor to compensate for classifier degeneracy and to improve the convergence of the wrapper search. The Q-factor, calculated from the ratio of sensitivity to specificity of the confusion matrix, is applied as a penalty to the accuracy (1-error rate). This method is successfully applied to a multiclass subject independent BCI using 10 untrained subjects performing 4 motor tasks in conjunction with the Sequential Floating Forward Selection feature search algorithm and Support Vector Machine classifiers.]

## 1. Introduction

Brain Computer Interface (BCI) is an ensemble of technologies that seek to establish a pathway of communication capable of translating neurologically derived signals, such as imagined human movements, into computer interpretable commands [1–3]. These commands can be used for the purpose of basic user interface, or for controlling external devices such as a robotic arm or prosthetic [1,4–8]. BCI implementation is a pattern recognition problem where signals derived from different brain states are examined in order to select a set of optimally descriptive features which can be most closely attributed to the associated state, typically using supervised learning.

Many successful approaches to BCI have been presented in the literature [2,9,10]. While many approaches focus on subject specific models due to high inter-subject variances present in larger populations, a wide array of studies have shown the viability of subject independent (SI) models [2,7,10–13]. Reducing complexity is an important part of emerging consumer grade EEG devices. Viability of an off the shelf subject independent solution relies on methods capable of being deployed for mobile devices and embedded platforms. While the computational capabilities of these devices are rapidly increasing, power requirements play a key role in feasibility of high computational complexity models. As a result, optimized models based on traditional machine learning techniques gain an advantage for BCI applications. Additionally, while BCI methods focus on interface and output classification, these models can be used to develop a more comprehensive understanding of the inner workings of the human brain for future research applications.

SIBCI methods aim to reduce cost and complexity through the selection of a set of features common across a large population. This approach is essential to the feasibility of real-world applications which require robust classification for a broad array of activities and events. Multiclass methods commonly employed in BCI literature include one-vs.-one (OVO) and one-vs.-rest (OVR) strategies which tend to generate highly comparable results [15–18]. In the interest of applying our method and findings to the widest available scale of both subject populations and event classifications, we conducted our investigation of multiclass detection using an OVR classifier scheme. As such, our selection of the OVR scheme with a data driven feature selection wrapper as described in section 2.2 allows us to optimize performance and reduce architectural complexity by selecting only one subset of features per class. To alleviate the undesirable behaviour sometimes present in classifiers operating on imbalanced sets, we have devised an OVR classifier performance metric to guide the feature search algorithm.

The Sequential Forward Floating Search (SFFS [19,20]) method was selected to operate in conjunction with the OVR multiclass Support Vector Machine (SVM) classifier to evaluate EEG data as processed by a variety of discrete and continuous output classifiers [21,22]. In this case the OVR approach scales better than OVO as the number of classes increases.

We will also show that OVR discrete output classifiers such as the SVM, when faced with features that are not well separated, can become trapped in degenerate states, where the classifier learns to always predict the dominant class. The impact of these degeneracies is increased when the classifiers are faced with imbalanced input classes. While metrics for performance measurement of BCI comprise only a small part of the active research related to the broader topic [23–28], application based concerns have drawn some interest to the investigation of imbalance resistant metrics [29–32]. Simple error rates do not produce a reliable measure when degenerate cases are encountered. Degeneracies are detrimental to the feature selection as searches can get trapped in local minima. A successful wrapper must detect and compensate for this condition.

In this paper we propose a novel scalar metric that indicates the successful detection rate and the degree of degeneracy of the classifier. We begin by providing descriptions of the multi-class SIBCI feature space, the degeneracies of the one versus the rest discrete output classifier, and the wrapper feature selection methods. We then provide an outline for our experimental design, feature selection methods and an overview of SFFS. This is followed by details of our proposed solution, and a presentation of our results which contains a comparison of corrected and non-corrected data processed by both subject independent and subject specific models. The final section provides an assessment of our results and a conclusion to the paper.

## 2. Background

### 2.1. BCI Feature Space

The inherent challenges of SIBCI are magnified in cases where feature sets are derived based on samples collected from larger populations, as these data sets tend to display substantial inter-subject variance. As it pertains to SIBCI, the inter-subject variance of electroencephalographic (EEG) signals [33] complicates the selection of task invariant features as more subjects are introduced. This principle can be illustrated for an arbitrary 2-dimensional feature space as shown in Fig 1. For a finite and fixed VC dimension, the classification error increases with the number of subjects or tasks [34,35].

**Fig 1.**
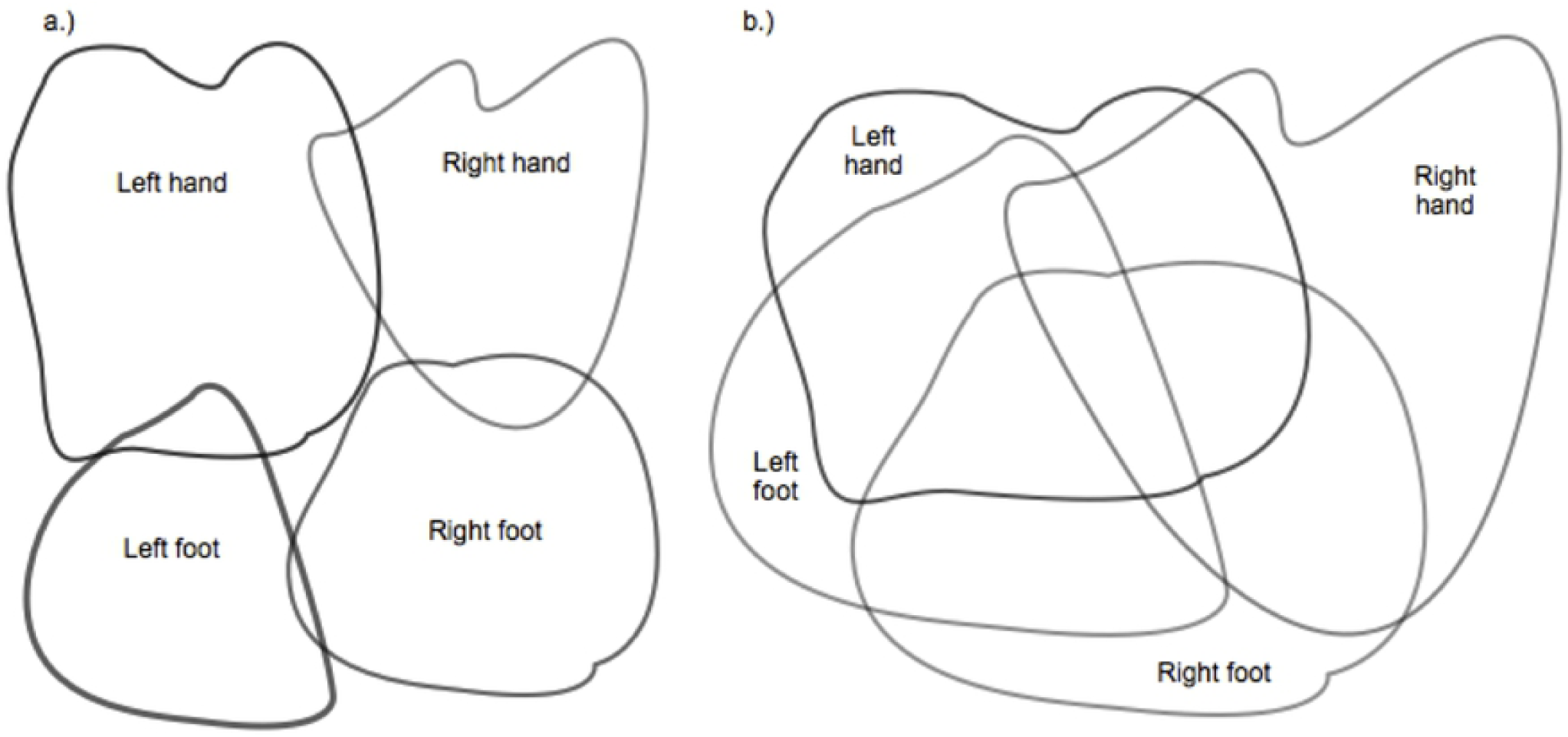
Conceptual illustration of multiclass BCI in feature space. (A) One subject with four tasks. (B) Multiple subjects with four tasks.

Representing each set of features for M tasks and P subjects as 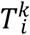, where *k* ∈ {1,2,3,…,M} corresponds to the number of tasks and *i* ∈ {1,2,3,…, P} the number of subjects, we describe the feature space for each task T^k^ as

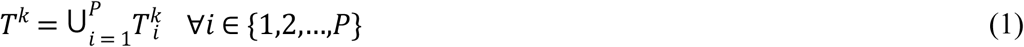

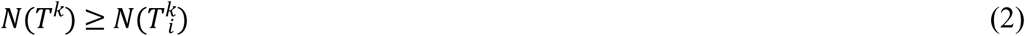

*N*(*T*) indicates the cardinality of each feature *T*, and indicates that the subject invariant task may occupy a larger area of the feature space than for any individual subject. Each *N(T)* increases as more subjects are included in T^k^. Fig 1A shows one possible feature space for *i* =1.

A unique feature space 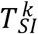 for each subject invariant feature *k* is required for correct operation of the classifier

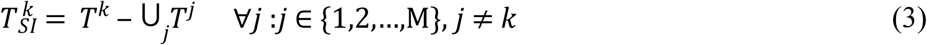

As M increases, the faster growth of the second term on the right-hand side reduces the probability of finding a location in the unique feature space for each individual task. This is presented graphically in Fig 1B. As additional subjects are included, the region uniquely occupied by each 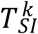 becomes smaller. The area of this region corresponds to the probability of correctly and uniquely mapping a feature to a task. This probability decreases as additional subjects or tasks are included (Blankertz et al. 2003).

Alternatively, Wolpaw et. al. [4] show this using the information capacity of a BCI channel, as derived from Shannon’s theorem [36,37]. Information transfer rate (ITR) is expressed in bits per second ITR = B/T, where B is the total sample length in bits and T is the sample duration. B shows the effect of increasing the number of tasks or classes on the probability of correct detection

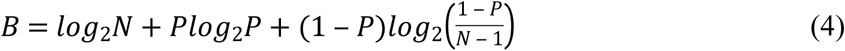

N equals the number of tasks and P is the probability of correct detection. The difficulty of increasing the number of tasks has been evidenced in the results of existing BCI research as few designs using more than 4 or 5 tasks have been successfully implemented [2,3,35,38].

### 2.2. Wrapper Methods

To obtain an efficient set of common features for subject SIBCI, we assembled a large number of high dimensionality feature vectors and conducted a non-exhaustive search for subsets of features to train SVM classifiers. A wrapper method chooses features by using an induction algorithm based on the performance of the actual classifier [39]. The search algorithm uses an optimization criterion derived from SVM feedback. As such, it is imperative that this optimization criterion accurately reflect the outcome of the classifier. Any degenerate mode that skews the results so as to ‘swamp out’ meaningful results will deteriorate the performance of the wrapper. This can result in wrapper algorithms that converge to incorrect solutions or fail to converge at all. Given an appropriate correction for this behavior, a wrapper method for feature selection can be well suited for multi-class SIBCI.

### 2.3. One versus the Rest classifier designs

Several approaches to the multiclass BCI classifiers can be found in the literature [3,9,10,35,40,41]. The OVO approach uses binary classifiers with each class containing an equal number of samples [15,16]. As M grows, a OVO solution to an M-class classification problem requires an exceedingly large M(M-1)/2 classifiers. Alternatively, the OVR approach treats the M class problem as M two-class problems thus requiring only M classifiers. A distinct feature subset is produced for each of the M classes or tasks. In contrast, the OVO arrangement increases the model complexity by requiring M(M-1)/2 feature selections. This results a factor of (M-1)/2 more feature subsets than tasks, none of which are uniquely assigned to a specific task. Additional methods are then required to select a single unique feature set for each task.

Generally speaking, for each class, there is an optimal discriminant function *g*_*i*_(***x***), *i =* 1,2,…, M, so that *g*_*i*_(***x***)> *g*_*j*_(***x***), ∀ *j* ≠ *i*, if *x* ∈ *ω*_*i*_. Designing the discriminant function so that *g*_*i*_(***x***)= 0 separates class *ω*_*i*_ from all of the others where each classifier should produce *g*_*i*_(***x***)>0 for *x* ∈ *ω*_*i*_, and *g*_*i*_(***x***)< 0 otherwise [42]. Classification is achieved according to the rule:

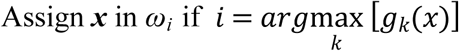

Indeterminate regions where either more than one or no *g*_*i*_(***x***) is positive can result from the overlapping regions such as those shown in Fig 1B.

An OVR classifier with an equal frequency of members in each of the M classes has M-1 greater number of training samples in the non-target class than in the target class. The unequal size of the two groups biases the overlapped region of Fig 1A toward the non-target class. The resulting classifier may fail to recognize members of the target class and impede the search algorithm.

Assuming equal number of samples per class, when an OVR classifier fails to recognize any members of its target class, the classifier accuracy is a deceptively high (M-1)/M (degenerate state). Examples of a perfect classifier, a typical classifier and two degenerate (null) classifiers are shown in Fig 2. In each example, we are looking at the outputs from discrete output classifiers. In the (M-1)/M degenerate state (Fig 2C), the classifier produces misleading accuracy of 75% for 4-classes.

**Fig 2.**
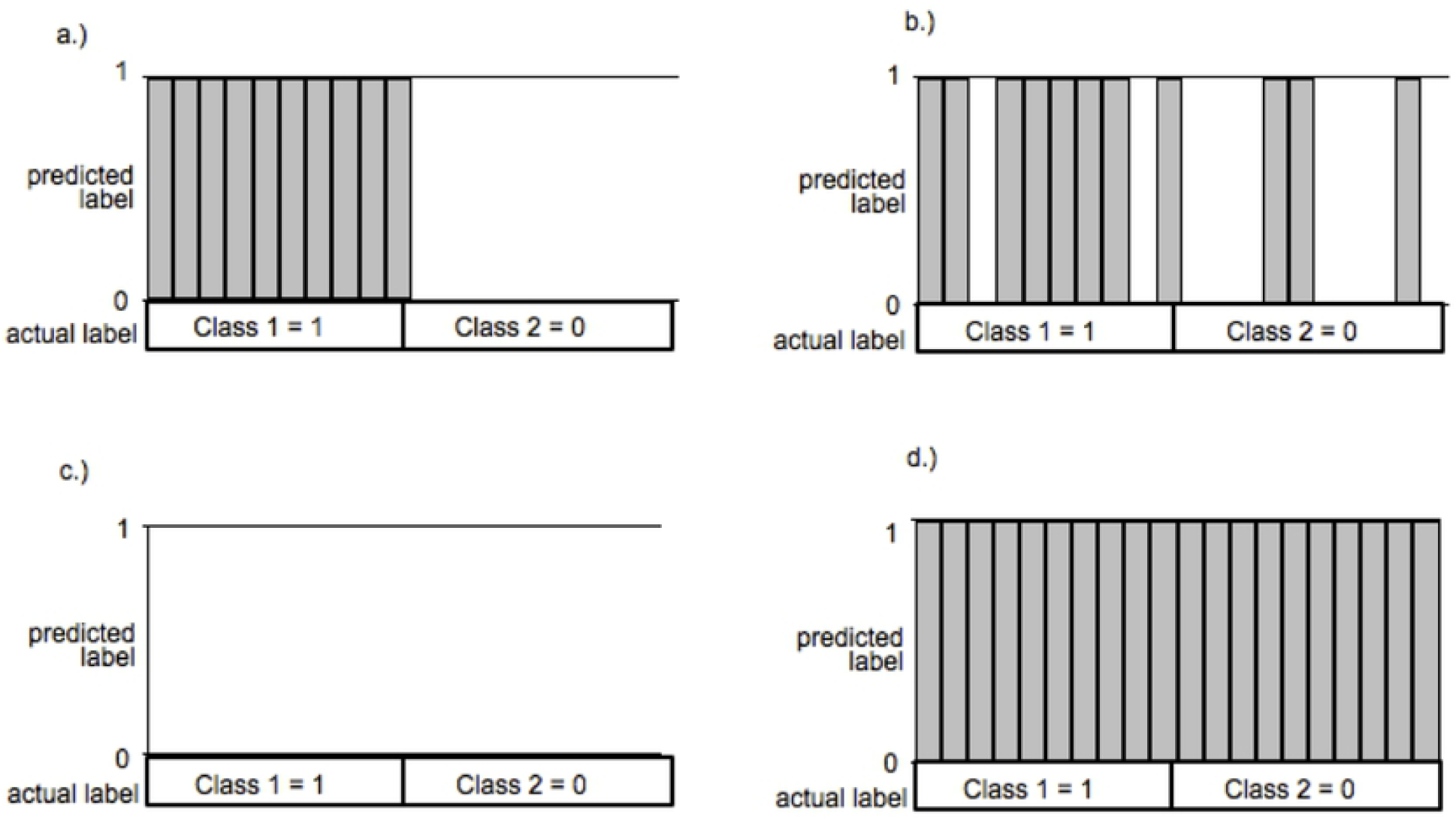
Plot of predicted vs. actual values for four classifiers examples. (A) Ideal classifier (B) Non-ideal classifier (C) Degenerate Classifier Mode I (D) Degenerate Classifier Mode II. Gray vertical bars indicate samples predicted to have a label of ‘1’. Lack of a gray vertical bar indicates a sample predicted to have a label of ‘0’.

Several metrics and methods have been demonstrated in the literature to evaluate performance of classifiers with imbalanced input classes. Among these metrics, we evaluated the G-mean [43], the F-measure [44], optimized precision [45], Cohen’s kappa coefficient [23,46], Jaccard’s coefficient [42], and balanced accuracy [47] for their suitability toward detection and mitigation of the degeneracy problem. Under-sampling and over-sampling methods were not considered as these only address the imbalance of the dataset, maybe to the detriment of learning, and do not provide a cost function metric. The behavior of each metric was analyzed for a small 4-class problem with 3 samples in each class. Each metric was evaluated to determine if it could provide a cost function that would consistently detect degeneracies within poorly performing classifiers without requiring additional scaling or thresholding.

The results are shown in Table 1.

**Table 1.** Classifier Metrics and Fitness values. Scalar metrics and fitness values are shown for 4 classes with 3 samples in each class. The designations for “Good” “Working” and “Poor” classifier are somewhat arbitrary and intended only to provide a relative comparison of similarly performing classifiers across each metric.

|  | Ideal | Good | Mode I | Mode II | Random | Moderate | Poor |
| --- | --- | --- | --- | --- | --- | --- | --- |
| True Pos | 4 | 3 | 0 | 4 | 1 | 2 | 1 |
| False Neg | 0 | 1 | 4 | 0 | 3 | 2 | 3 |
| False Pos | 0 | 3 | 0 | 12 | 9 | 4 | 4 |
| True Neg | 12 | 9 | 4 | 0 | 3 | 8 | 8 |
| accuracy | 1.0 | 0.75 | 0.5 | 0.25 | 0.25 | 0.63 | 0.56 |
| sensitivity | 1.0 | 0.75 | 0.0 | 1.0 | 0.25 | 0.50 | 0.25 |
| specificity | 1.0 | 0.75 | 1.0 | 0.0 | 0.25 | 0.67 | 0.67 |
| precision | 1.0 | 0.50 | undef. | 0.25 | 0.10 | 0.33 | 0.20 |
| recall | 1.0 | 0.75 | 0.0 | 1.0 | 0.25 | 0.50 | 0.25 |
| G-mean | 1.0 | 0.75 | 0.0 | 0.0 | 0.25 | 0.58 | 0.41 |
| F-meas. | 1.0 | 0.60 | 0.0 | 0.40 | 0.14 | 0.40 | 0.22 |
| Q-factor | 1.0 | 0.75 | 0.0 | 0.0 | 0.25 | 0.47 | 0.21 |
| Opt. Prec. | 1.0 | 0.75 | -0.75 | -0.25 | 0.25 | 0.40 | -0.10 |
| Bal. acc. | 1.0 | 0.75 | 0.50 | 0.50 | 0.25 | 0.58 | 0.46 |
| Cohen's | 1.0 | 0.43 | 0.0 | 0.0 | -0.33 | 0.14 | -0.08 |
| Jaccard's | 1.0 | 0.43 | 0.0 | 0.25 | 0.08 | 0.25 | 0.13 |

## 3. Experimental Design

A four-class SIBCI was implemented using scalp level EEG signals from 11 electrodes from the pre-frontal, pre-motor and motor cortex regions. Electrode locations Fp1, Fp2, F3, F4, F7, F8, T3, T4, C3, C4 and Cz from the pre-frontal and pre-motor cortex regions were included because of their role in planning of imagined motor tasks [48]. Our target classes consisted of left hand, right hand, left foot, and right foot imagined movements based on our own research [22] and that found in the literature [2,3,9,38,41,49]. The signal processing stages include feature extraction, feature selection, and classification.

### 3.1 Materials and Methods

Study protocol include the enrolment of ten untrained volunteers who provided written verification of their informed consent to participate. Protocols employed in the study were conducted under the supervisory authority of the University of Missouri Kansas City Institutional Review Board under protocol #090218. Neurological data were collected using standard 10-20 electrode placement with two ear electrode references. Signals were amplified and digitized using a *NeuroPulse Systems* MS-24R *bioamplifier* and 1.5 – 34 Hz bandpass filter with a 256 Hz sampling rate.

Each volunteer was asked to perform four incremental or quasi-movements [21,22,50] without rehearsal for 8 seconds. Two identical sessions were conducted 4-6 weeks apart, each consisting of 12 repetitions of the four intended movements.

Each volunteer was visually prompted to produce a restrained hand or foot movement while gripping a rubber ball in each hand or pressing the appropriate foot into a rubber floor mat. The initiation of each visual prompt was accompanied by a short 500 Hz aural alert cue beep. The volunteers were requested not to blink prior to and during the performance of the quasi-movement. A 10 second relaxation period was provided between each series of movements. Only the first 2 seconds of each movement event were analyzed.

As a result, 960 8-second signal segments were collected. The first and last repetitions of each session were discarded, leaving 800 segments for processing and feature extraction. Of these 800 segments, 200 were left out for testing and 400 were used for training and 200 for cross validation in accordance with a 3-fold validation.

### 3.2. Feature Description

After collecting, filtering and segmenting the EEG signals, we implemented an extraction method focusing on the eight feature types. These eight features were selected for their descriptive power and relative ease of generation based on methods which we had successfully employed in previous investigations [21,22].

#### 3.2.1. Linear Predictive Coefficients

Linear predictive coding (LPC) [51] filter coefficients represent the time domain behavior of complex linear signals and have been successfully used for EEG processing and BCI applications [52,53]. After applying a 3x downsampling, coefficients of an 18^th^ order LPC (parametrically selected based on previous empirical observations) were chosen as features.

#### 3.2.2. Power Spectral Density

Event-related desynchronization of neuronal signals as a result of imagined motor task have been reported in the literature [54,55]. This desynchronization has been observed as changes in the amplitude of power spectral density (PSD) energy in the EEG signals. By using Welch periodograms and Hamming window length of 33 with 97% overlap, spectral amplitude coefficients were generated. The coefficients representing frequency components from 1 - 48 Hz were aggregated into twelve 4 Hz frequency bins. The spectral energy in each bin was calculated and the coefficients were used as PSD features.

#### 3.2.3. Cepstrum

Coefficients of the Cepstrum, a non-linear Fourier Transform of the EEG spectrum, were collected and used as features. Cepstrum was included for its possibility to detect nonlinear features [56].

#### 3.2.4. Short Time Fourier Transform

Short time Fourier transforms (STFT) were used to capture the non-stationary properties exhibited in the time-frequency behaviour of EEG signals [55]. STFT was extracted on the down-sampled EEG signals using a sliding window length of 1 second and an 84 sample overlap. The signal amplitude coefficients with the frequency span of 1 – 48 Hz were aggregated into four 12 Hz frequency bins and the energies under each bin were used as features. These parameters were selected by trial and error.

#### 3.2.5. Wavelets

Wavelet decomposition coefficients from filter banks and wavelet packets were included as features to capitalize on the multi resolution capabilities of their spectro-temporal modalities [57]. Filter bank coefficients and wavelet packets produced two sets of features. Another set of features was generated from energies marginalized over shift (time). These energies were calculated over different scales (frequencies). In our previous studies [21,22] the classification performance of biorthogonal spline, reverse biorthogonal spline, Morlet, Coiflet, symlet, Gaussian, Daubechies, Meyer wavelets and their variants were examined. Based on the results of these studies, we selected reverse biorthogonal spline 3.7 wavelets for wavelet decomposition filter bank coefficients, symlet 15 wavelets for wavelet packet decomposition filter bank coefficients, and reverse biorthogonal spline 3.1 wavelets for both sets of energy features.

#### 3.2.6. Common Spatial Pattern (CSP)

Common Spatial Pattern Filtering (CSP) of the EEG signals was used to generate additional sets of features. EEG CSP filtering has been extensively used in BCI [41,54,58,59]. This method is especially useful for the detection of event related desynchronization [58].

For our multiclass problem we projected each EEG signal segment, **E**, into a CSP decomposition, **Z**, according to a CSP projection matrix, **W** according to

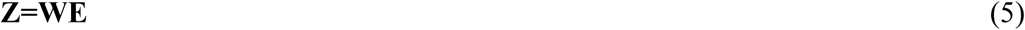

The elements of **W**^-1^ represent common spatial patterns (i.e, time invariant EEG source distributions) [58].

### 3.3. Feature Selection

The objective of feature selection is to find a salient subset of features that leads to the smallest classification error [19,60]. To this end, we chose Sequential Forward Floating Selection (SFFS) within our wrapper method. SFFS avoids the nesting problems of the Sequential Forward Selection method, where once a feature is chosen it cannot be discarded; and a similar issue with the Sequential Backward Selection, where a feature cannot be included in the final subset once it has been discarded [20,42]. SFFS re-evaluates previously discarded features for inclusion while previously selected features are also re-evaluated for discard. A detailed treatment of SFFS is provided by Theodoridis [42]. In general, we consider a set of *m* features and build the best subset *k* for *k = 1, 2*, …, *l* ≤ *m* where a cost criterion C is optimized. Let X_k_ = {x_1_, x_2_, …, x_k_} be the set of the best combination of features, and Y_m-k_ represent the remaining unselected *m – k* features. Retaining all of the remaining lower dimension subsets X_2_, X_3_, …, X_k-1_ we build on the sets by creating the next subset X_k+1_ by choosing an additional element from Y_m-k_. The newly created features are compared with the original subsets and if the cost function value is improved, the new feature is retained. The array of costs C_2_, C_3_, …, C_k+1_ associated with X_2_, X_3_, …, X_k+1_ may assume a minimum value that leads the algorithm to pick a feature subset resulting in a degenerate classifier, or the SFFS search may fail to converge.

### 3.4. Support Vector Machine Classifiers

SVMs are kernel based maximum margin classifiers [61,62]. We examined different Gaussian kernels, with spread values σ ranging from 1 to 100 and box constraint or C-parameter values of .001 to 1000 to find the best parameters. Smaller values of C reflect soft margins allowing the SVM to overlook outliers, while larger values indicate hard margins, resulting in a rigid and possibly over-trained classifier. [34,49,63].

## 4. Addressing Degeneracy

### 4.1. Degeneracy Detection

Detection of degenerate and non-degenerate classifiers can be accomplished by using elements of the confusion matrices of the classifier because for discrete output classifiers, ROC AUC methods cannot be readily applied, as the output produces a single point in ROC space [64]. A confusion matrix **A** = [*A(i, j)*] is obtained from tabulating the number of data points with true class *i* designated as class *j* by the classifier. Hence, for an OVR classifier, unfortunately, since the confusion matrix is not a scalar value, multiple confusion matrices cannot be easily compared when assessing classifiers. For this reason, a comparison of the Sensitivity and Specificity is helpful. For an OVR classifier with a confusion matrix **A** ∈ ℝ^2 × 2^ we desire a transformation into a scalar value which preserves both the accuracy information while accounting for possible degeneracies in the solution. To achieve this, we devised a metric henceforth dubbed as the Q –factor which is defined as:

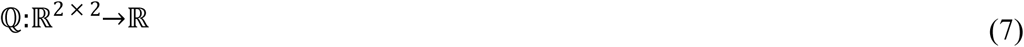

From elements of **A**, we can obtain overall accuracy (ACC), Sensitivity (SENS) and Specificity (SPEC) as follows:

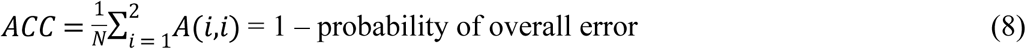

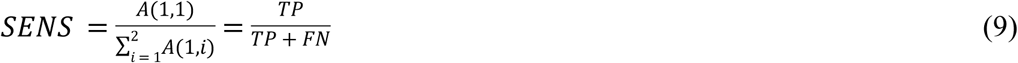

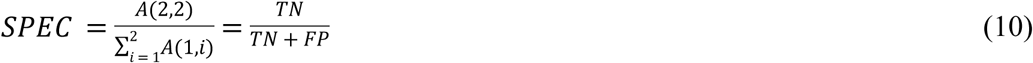

Naturally constrained by the scale of total observations:

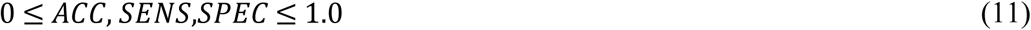

A full summary of the possible classifier states for the classifiers shown in Fig 2 (with M total number of classes) is given in the Table 2.

**Table 2.** States and performance metrics of possible classifiers.

| Parameters | Classifier Type | Ratio |
| --- | --- | --- |
| $ACC = SPEC = SENS = 1.0$ | ideal | $SENS/SPEC = 1$ |
| $0 < ACC \approx SPEC \approx SENS < 1.0$ | Non-ideal | $SENS/SPEC < 1$ |
| $SENS = 0; SPEC = 1.0; ACC = \frac{M-1}{M}$ | Mode I degenerate | $SENS/SPEC = 0$ |
| $SENS = 1.0; SPEC = 0; ACC = \frac{1}{M}$ | Mode II degenerate | $SPEC/SENS = 0$ |

We designate a trivial classifier that always rejects instances of the target class as Mode I degenerate. Conversely, we designate a classifier that always accepts such instances as Mode II degenerate.

Mode I is the most detrimental to an ACC-driven wrapper search since 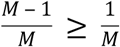 for all possible values of M.

The SPEC/SENS ratio indicates the imbalance in the off-diagonal elements of the confusion matrix **A**, and thus quantifies the amount of degeneracy in the classifier.

The Imbalance ratio

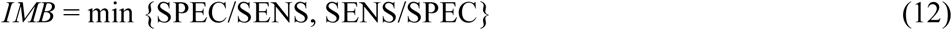

is the minimum of either the SPEC/SENS or its inverse. This detects both modes while eliminating any infinite values. We now transform the confusion matrix to a scalar via F(x,y) where x = *IMB* and y = ACC.

From Table 1, we can see that geometric mean (G-mean), Q-factor, optimized precision and Cohen’s K factor each assign either zero or negative fitness values to mode I and mode II classifiers. Only G-mean and Q-factor assume a minimum of zero when facing poorly functioning classifiers. Further, the Q factor produces a greater penalty for the poorly performing classifiers when compared with G-mean.

### 4.2. Degeneracy Correction

If a wrapper method selects features considering the imbalance in the confusion matrix in addition to simple fitness metrics such ACC, it may avoid degenerate states exemplified in Fig 2C-D. Fig 3A further visualizes ACC values vs. confusion matrix imbalance. Dividing the accuracy vs. imbalance space into 4 separate regions reveals region 1 of high relevance and high validity, where the most desirable results occur. Region 2 of high relevance and low validity are where the most detrimental results occur. Regions 3 and 4 are low relevance. The most desirable results on the plot are located along the upper left boundary of Region 1 along the vertical axis.

**Fig 3.**
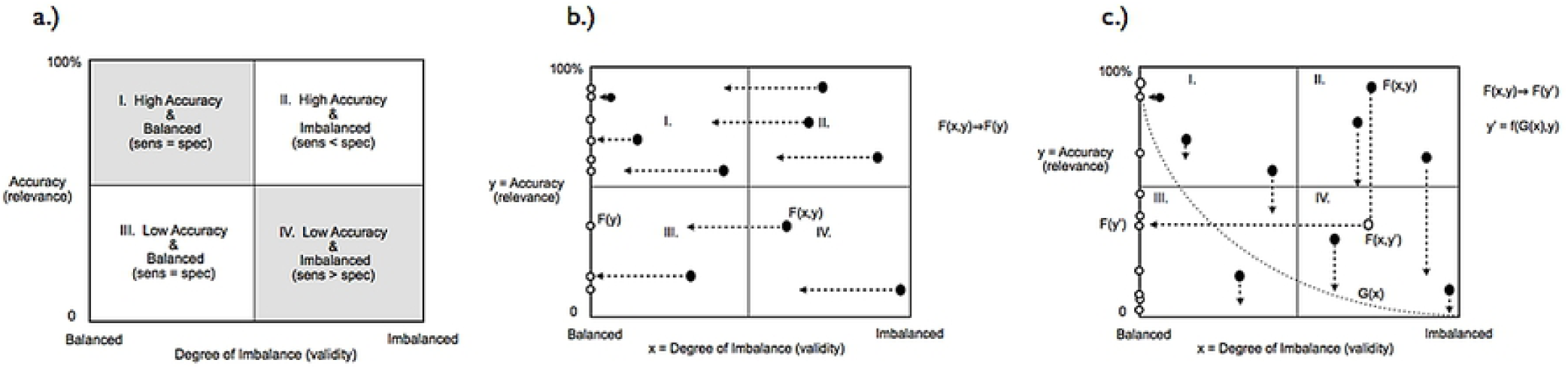
Conceptual plots of confusion matrix imbalance vs. accuracy. (A) Highlighting the 4 regions of High and Low Relevance and High and Low Validity (B) Projection of features onto the Accuracy axis without considering imbalance(C) Projection of features onto the Accuracy axis after application of the Q-factor. These plots are illustrative only and do not represent actual data. Actual data plots are shown in Fig 5.

Thus we can better assess our features by confusion matrix’s imbalance in addition to accuracy. While a simple accuracy calculation (Fig 3B) transforms the information to a one dimension scalar value as required by the wrapper, it fails degenerate cases with high imbalance.

### 4.3 Q-factor Derivation

We propose a simple yet effective conversion of the classifier accuracy metrics into a scalar value to guide wrapper methods and to avoid degenerate states. T aforesaid scalar, dubbed the *Q-factor*, is defined as

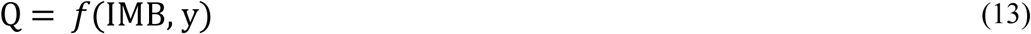

Where y is the simple accuracy, and IMB, the degree of imbalance, is given by (12). This accomplishes our objective of moving any F(x,y) with a large value of imbalance into a region of lower relevance and imbalance (Fig 3), clearing out the elements located in Region 2 and moving them into Regions 1, 3 and 4 as shown in Fig 3C.

Finally, the Q-factor is given by

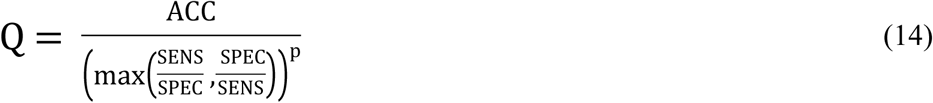

*p* is used to adjust the imbalance behavior and is nominally set to *p* = 1. A larger value of *p* will move the boundary between Region 1 and Region 2 closer to the left vertical axis.

### 4.4 Application to multiclass BCI

Fig 4 depicts the wrapper problem for a hypothetical 6-class problem to illustrate the effect of Mode I degeneracy. The increasing logarithmic curvature of the upper line describes the locus of the Mode I degeneracy state as described in Section 2.2. The Mode II line corresponds to the 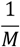 Mode II degeneracy state. The dashed line represents the constant bit rate line for an estimated value of our SIBCI from *M* = 2 to *M* = 10, based on Eq. (1). For a fixed bit rate channel, the limit of the probability of a correct decision decreases as the number of classes increase. For an OVR classifier, the impact of Mode I degenerate results becomes more severe as M increases. If the degenerate result is near or greater than the theoretical limit for a given class, the wrapper has a greater chance of basing its search on the degenerate results.

**Fig 4.**
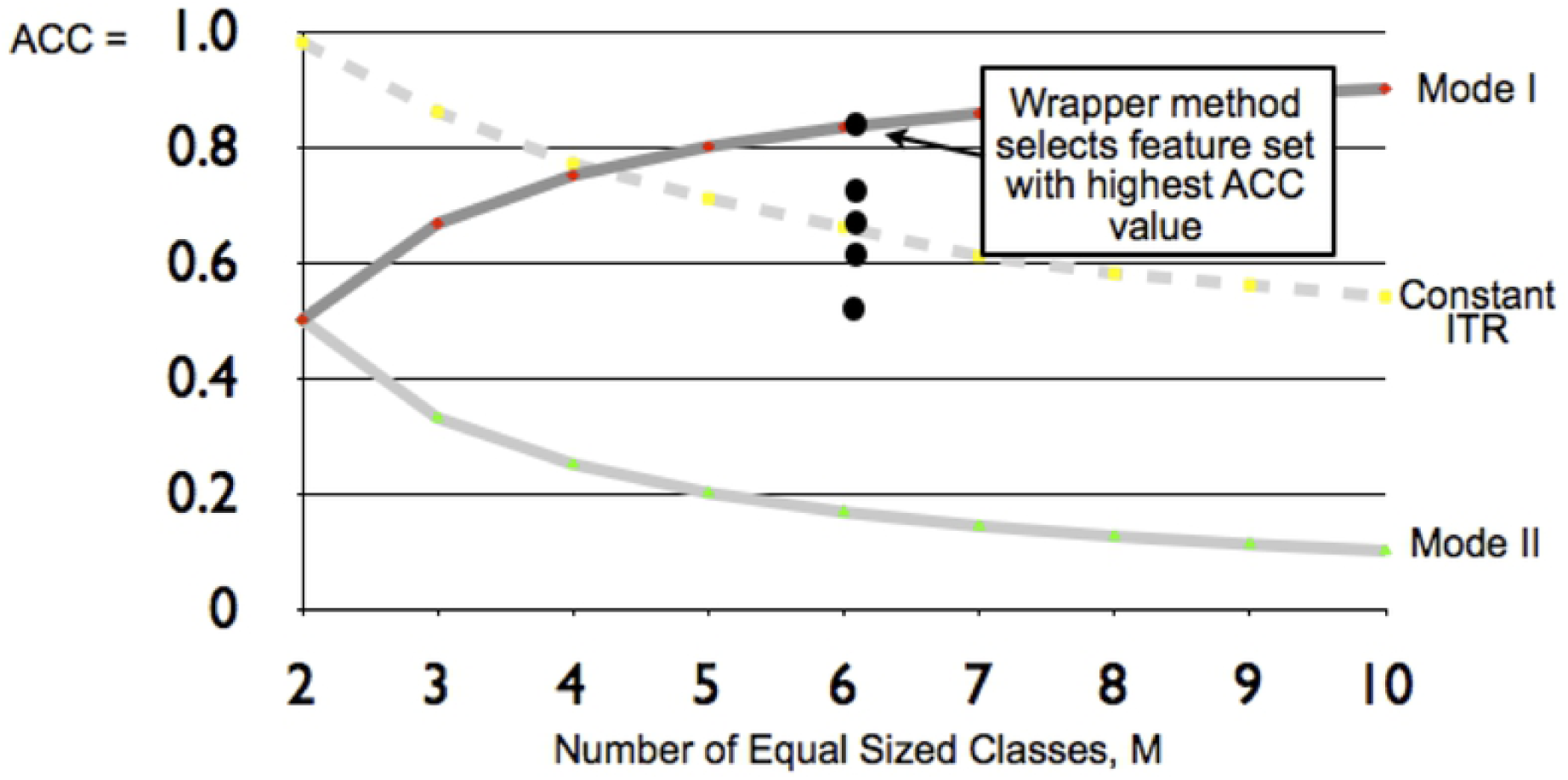
Plot showing effect of Mode I degeneracy for one possible 6-class problem. Accuracy vs. number of classes is shown for Mode I and Mode II degenerate states. Dashed line indicates the calculated accuracy limits according to the bit per trial Information Transfer Rate.

## 5. Results

For comparison, we conducted both Q-factor guided and simple accuracy guided feature searches on subject specific BCI (SSBCI) classifiers using our earlier-mentioned 10 subject BCI dataset (Fig 5A-B). The results of that comparison are presented in Table 2. After application of the Q factor to the classifier, the SSBCI results moved out of region 2 (high accuracy - high imbalance) into regions 1, 3, and 4. Prior to applying Q to our model, we found that the SFFS search algorithm more frequently became trapped in local minima and was therefore unable to produce a viable feature subset. The total number of convergent SFFS searches that produce feature subsets is indicated in Table 3.

**Table 3.** Summary of Results. Mean and standard deviation values for Subject Independent (SI) and Subject Specific (SS) BCI Features before and after Q-factor corrections. Imbalance is calculated by 1-min{sens/spec, spec/sens} where 1 is worst and 0 is the best. CSP indicates common spatial pattern features.

|  | Imbalance |  | Accuracy (%) |  | Number of<br>Non-degenerate<br>Classifiers |
| --- | --- | --- | --- | --- | --- |
|  | Mean | Std. Dev. | Mean | St. Dev. |  |
| Uncorrected (ACC) |  |  |  |  |  |
| SI | 0.7051 | 0.1930 | 72.75 | 40.48 | 177 |
| SS | 0.2991 | 0.2829 | 89.44 | 8.64 | 18089 |
| SI CSP | 0.0707 | 0.636 | 56.78 | 9.70 | 221 |
| Corrected (ACC * Q-factor) |  |  |  |  |  |
| SI | 0.1635 | 0.2695 | 64.41 | 7.30 | 3719 |
| SS | 0.0384 | 0.0958 | 80.34 | 12.82 | 20121 |
| SI CSP | 0.0920 | 0.1794 | 56.94 | 11.26 | 1798 |

**Fig 5.**
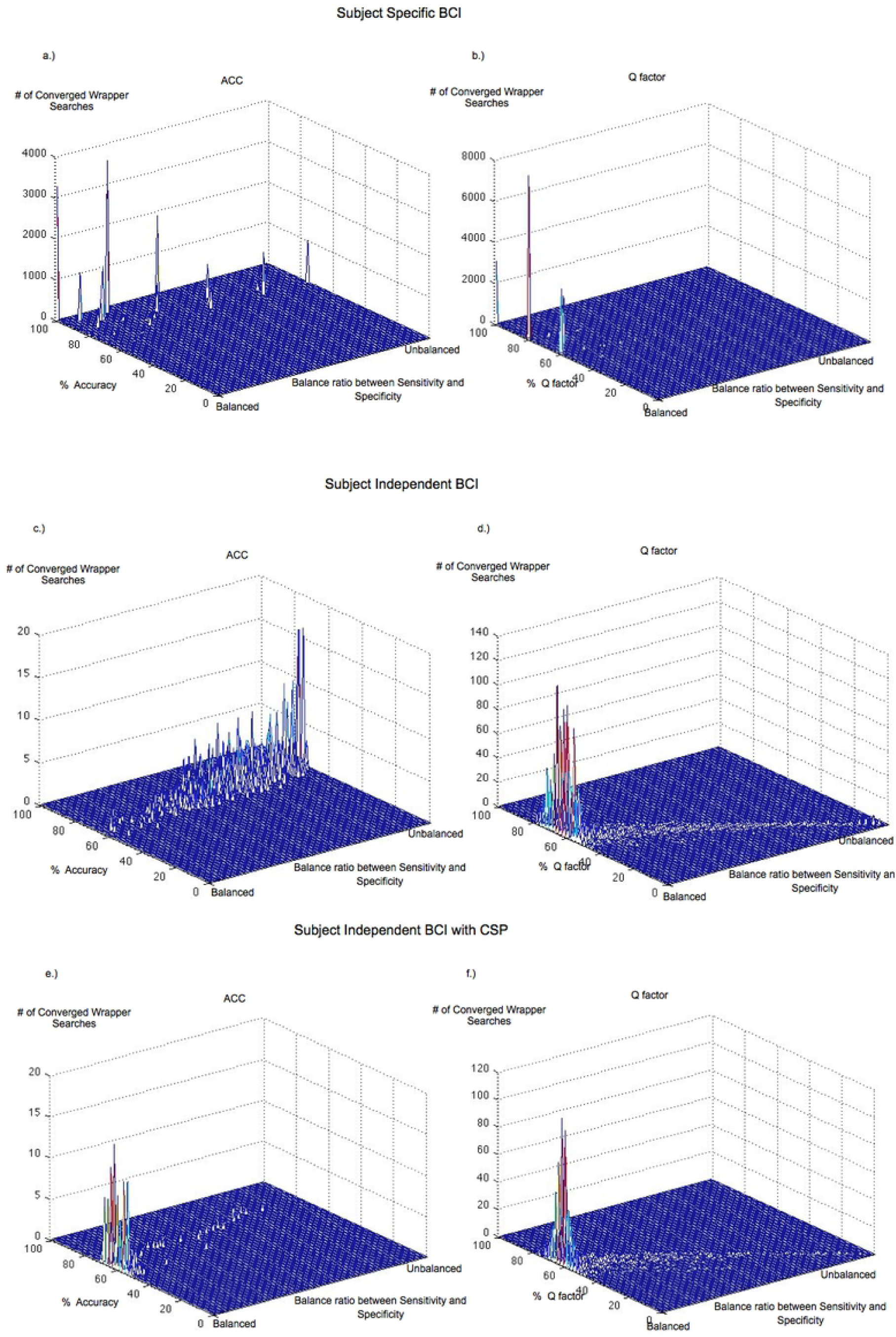
Accuracy vs. imbalance space for our 10 subject, 4 movements experiment (all modalities, cross validation results). (A) SSBCI before Q-factor correction (B) SSBCI after Q-factor correction (C) SIBCI before Q-factor correction (D) SIBCI after Q-factor correction (E) SIBCI with CSP before Q-factor correction (F) SIBCI with CSP after Q-factor correction

After incorporation of the Q factor an increase in converged searches was produced. Q-corrected results occur tightly clustered to the vertical axis in region 1. This is the optimal location for classifier accuracy-imbalance results, indicating the highest degree of valid results due to equal sensitivity and specificity of the confusion matrices.

Comparing the Subject Independent BCI (SIBCI) results after applying the Q-factor (Fig 5D) to the uncorrected SIBCI (Fig 5C, we observe that upon applying the Q factor to our SIBCI wrapper searchers, two improvements are achieved. First, invalid results in region 2 are cleared out and moved to Regions 1, 3 and 4. Second, the number of converged wrapper searches are increased. This is attributed to the higher quality of Q-factor feedback to the SFFS algorithm with its integrated SVM in the wrapper. Degenerate outputs were downgraded by the Q-factor and did not guide the search. Most classifier results are concentrated along the vertical axis of region 1. Q-factor corrected results occurring near the vertical axis, 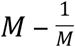, are still located near the Mode I degeneracy limit (75%), however the results were based on working classifiers and are not due to the inclusion of a large number of degenerate classifiers.

When applying the Q factor to the CSP model (Figs. 5e & 5f), few features or classifier outputs occurred in Region 2. A greater number of converging wrapper models were produced, and upon applying the Q factor they were tightly clustered against the vertical axis of region 1. The summary of results in Table 3 shows three areas where Q factor improves the performance.

First, the number of SFFS searches that produce feature sets has increased in all cases, and more so for SIBCI models. While a small increase in the number of converged wrapper models occurred for the SSBCI, the improvement is not as pronounced. Uncorrected SIBCI wrapper searches encounter more difficulty in convergence due to the lower discrimination of features as described earlier, and thus show larger improvements when using Q-factor.

Second, we observe that the imbalance for all models improves after the application of the Q factor. The improvement is most pronounced for SIBCI than for SSBCI, except for SIBCI with CSP. We conjecture this is due to uncorrected CSP models not exhibiting a high degree of imbalance in the first place.

Third, in most cases, the standard deviations of the Q-corrected accuracies and imbalances for SIBCI are less than those of the uncorrected values. All corrected SIBCI accuracy and imbalance results are tightly clustered around their respective mean values. The means of Q-corrected accuracy values are slightly lower than the uncorrected values (73% vs. 64% for SIBCI). This is attributed to the Mode I degeneracy producing an accuracy value of 75% for M=4 (using M-1/M). The reduction in accuracy is caused by the elimination of this bias.

Finally, features that occurred most frequently in our Q-corrected wrapper output were power spectral density, Cepstrum, short-time Fourier transform and wavelet decomposition energies marginalized over time.

## 6. Conclusion

This study was based on our in-house SIBCI dataset using cross-validation to mitigate the smaller size of the dataset. The Q factor was successful in eliminating trivial (null) and random classifiers of the wrapper feature selection method. As a result, we found a larger number of suitable feature sets. Subject independent results showed a high degree of imbalance when left uncorrected without Q-factor. We showed that while our Q-factor yields some improvements for SSBCI and CSP filtered SIBCI, the improvements were more pronounced for SIBCI. This stands to reason since SIBCI feature sets are more difficult to separate and produce a larger number of imbalanced classifiers within the wrapper search given the complexity of subject-the independent case.

Our uncorrected four class SIBCI results approach the 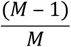 Mode I degenerate accuracy value of 75%. After applying the Q factor, SIBCI indicated less influence from the Mode I degeneracy.

Several topics have been identified to extend the usefulness of this method in future studies. Other mathematical, parameter variations, or transformation methods were not explored and have been deferred for future work. Experiments with M>4 classes is deferred to future research.

## 7. Acknowledgements

This work was supported in part by a University of Missouri Research Board and University of Missouri – Kansas City Faculty Research Grant.

